# Induced pluripotent stem cell-derived primary proprioceptive neurons as Friedreich ataxia cell model

**DOI:** 10.1101/829358

**Authors:** Chiara Dionisi, Myriam Rai, Marine Chazalon, Serge N. Schiffmann, Massimo Pandolfo

## Abstract

Human induced pluripotent stem cells (iPSCs) are used to generate models of human diseases that recapitulate the pathogenic process as it occurs in affected cells. Many differentiated cell types can currently be obtained from iPSCs, but no validated protocol is yet available to specifically generate primary proprioceptive neurons. Proprioceptors are affected in a number of genetic and acquired diseases, including Friedreich ataxia (FRDA).

FRDA is a recessive neurodegenerative and systemic disease due to epigenetic suppression of frataxin (FXN) expression caused by the presence of expanded GAA repeats at the *FXN* locus. The most characteristic early neuropathologic finding in FRDA is the loss of large primary proprioceptive neurons in the dorsal root ganglia (DRGs), with associated loss of large myelinated fibers in the dorsal roots and in the posterior columns of the spinal cord. Both a developmental deficit and progressive neurodegeneration are thought to underlie the loss of proprioceptors in FRDA, though the relative contribution of these two components is unclear. The basis of the high specific vulnerability of proprioceptors in FRDA is also unknown. In order to address these open questions about FRDA pathogenesis and at the same time develop a cell model that can be applied to other conditions primarily affecting proprioceptors, we set up a protocol to differentiate iPSCs into primary proprioceptive neurons. We modified the dual-SMAD inhibition/WNT activation protocol, previously used to generate nociceptor-enriched cultures of primary sensory neurons from iPSCs, to favor instead the generation of proprioceptors. We succeeded in substantially enriching iPSC-derived primary sensory neuron cultures in proprioceptors, largely exceeding the proportion normally represented by these cells in dorsal root ganglia. We also showed that almost pure populations of proprioceptors can be purified from these cultures by fluorescence-activated cell sorting. Finally, we demonstrated that iPSCs from a FRDA patient can generate normal appearing proprioceptors but have subtle differentiation deficits and more limited survival.

## Introduction

Human induced pluripotent stem cells (iPSCs) have been used for a decade to generate models of human diseases that recapitulate the pathogenic process as it occurs in affected cell types. Though the context of a cell culture dish is obviously different from the living animal or human being, many cell-autonomous processes can be reconstructed and analyzed in hiPSC-derived cultures, as well as interactions among different cell types that can be co-differentiated or co-cultured, even in 3D^1^.

Friedreich ataxia (FRDA) is an autosomal recessive multisystem disorder with prominent neurological manifestations, often associated with hypertrophic cardiomyopathy, skeletal abnormalities and carbohydrate intolerance. In almost all cases, the underlying genetic mutation is the hyperexpansion of a GAA repeat in the first intron of the *FXN* gene, encoding frataxin (FXN). The expanded GAA repeats promote chromatin condensation and disrupt *FXN* mRNA transcription^2^, leading to markedly reduced FXN levels in affected individuals. Reduced FXN levels impair the mitochondrial biogenesis of iron-sulfur (Fe/S) clusters^3^, leading to loss of activity of multiple Fe/S enzymes, mitochondrial dysfunction, and altered iron metabolism. However, despite notable progress, neither the details of these pathogenic processes nor the reasons why most cells do not seem to be functionally impaired by low FXN levels while specific cell types are vulnerable^4^ are yet fully understood. The earliest neuropathological finding in FRDA is the loss of large, parvalbumin-positive primary sensory neurons in the dorsal root ganglia (DRGs), corresponding to proprioceptors, and the associated loss of large myelinated fibers in the dorsal roots and in the posterior columns of the spinal cord^5^. This pathology leads to the onset and initial progression of ataxia and to the characteristic loss of deep tendon reflexes already found in most FRDA patients at the time of diagnosis. Cerebellar and pyramidal degeneration are then mostly responsible for disease progression^4^. It is also unclear to what extent the pathology in the proprioceptive system in FRDA is developmental or degenerative. Neuropathology and neurophysiology studies showed that DRG and spinal cord pathology is scarcely progressive, suggesting that it may be at least partly developmental^6, 7^. However, DRGs from patients with long disease duration still show signs of active invasion of large primary sensory neurons by satellite cells and inflammation, supporting a continuing degenerative process^5^. The question is critical for translational research in FRDA. We^8–10^ and others^11–13^ have characterized the genetic, epigenetic and biochemical phenotype of iPSC-derived “default” neurons, close to immature cortical neurons, showing that they recapitulate essential aspects of FRDA pathogenesis that are at least partially corrected by restoring FXN levels, but no FRDA cell model consisting of human proprioceptors is yet available. Although in some studies iPSC neural differentiation has been cued toward a neural crest-primary sensory neuron fate^14^, or even to primary sensory neurons^11^, they did not specifically enrich for or purified proprioceptors. A rodent cell model consisting in DRG cultures from conditional knock-out mice whose *Fxn* gene was deleted in Parvalbumin-positive neurons, including proprioceptors, has also been developed, but it has limitations such as the lack of expanded GAA repeats and the post-development induction of FXN deficiency. For these reasons we developed a protocol allowing to differentiate iPSCs into DRG-like cultures that are highly enriched in proprioceptors, which can be further purified by fluorescence-activated flow sorting (FACS). Obtaining such cultures required mimicking the process of DRG differentiation in a culture dish, while enhancing the signaling and neurotrophic processes supporting the differentiation of proprioceptive neurons.

DRGs contain a heterogeneous population of primary sensory neurons relaying temperature, pressure, pain and body position and movement (proprioception) to the central nervous system. These neurons differ in cell body size, axon diameter, degree of myelination, peripheral targets and efferent terminals in the spinal cord. Small diameter nociceptors have thinly myelinated or unmyelinated axons and express the neurotrophin receptor TrkA, one of the members of the tropomyosin-receptor kinase (Trk) subfamily of tyrosine kinases receptors, that has high affinity for Nerve Growth Factor (NGF).

Mechanoreceptors have large diameter fibers and detect mechanical stimuli via specialized end organs localized in the skin, such as Meissner’s corpuscles, Pacinian corpuscles, Ruffini corpuscles and Merkel cell endings. They express the neurotrophic receptor TrkB, which has high affinity for Brain Derived Neurotrophic Factor (BDNF). Some mechanoreceptors also express the neurotrophin receptor TrkC, activated by Neurotrophin-3 (NT-3). Proprioceptors represent about 7.5% of DRG neurons and have the largest cell bodies^15^. They express the Ca^2+^ binding protein parvalbumin^16^ and, at least in the mouse, high levels of FXN, accounting for most FXN found in DRGs^17^, a possible clue to their specific vulnerability in FRDA. Proprioceptor subpopulations can be identified by the expression of additional markers such as CDH13 and CRTAC1^18^. They have large (12-20 µm) and medium (6-12 µm) diameter myelinated axons (Aα, Aβ fibers) with fast conduction velocity (72-120 m/s for large and 30-70 m/s for medium proprioceptors). Specialized proprioceptive end organs include muscle spindles and Golgi tendon organs. The central branches of their axons climb to the medulla forming the posterior columns of the spinal cord and send branches to Clarke’s column, to laminae III-V of the spinal cord, and directly to motor neurons. These connections are the anatomical substrate of conscious proprioception, provide proprioceptive information to the cerebellum via the spinocerebellar tracts, and underlie the monosynaptic stretch reflex.

Primary sensory neurons in DRGs derive from a population of neural crest cells, induced by BMP, TGF-β and WNT signaling^19^, that is specified at the border between neural and non-neural ectoderm in the dorsal neural tube during gastrulation, then delaminates from the neural tube and migrates along a ventral pathway^20^. Coupled to migration, sensory neurogenesis occurs in two successive waves. The first wave is guided by the transcription factor Neurogenin 2^21^ and preferentially generates TrkB^+^ and/or TrkC^+^ mechanoreceptors and proprioceptors. The second wave is guided by Neurogenin 1^21^ and gives rise to small TrkA^+^ nociceptive neurons, but also to large TrkB^+^/TrkC^+^ neurons, as corroborated by the finding that in Neurogenin 2-null mice Neurogenin 1 is sufficient for these neurons to develop normally. Neurogenins initiate pan-neuronal programs but do not specify neuronal subtypes in the sensory lineage. This step requires the action of the Trk receptors and their cognate neurotrophins, which control cell survival, axonal growth, target innervation and the establishment of functional contacts between cells, and also regulate peptide and ion channel expression^22^. The RUNX family of transcription factors are key intrinsic mediators of these processes. RUNX1 and RUNX3, are expressed in DRGs after the neurogenins and act non-redundantly to promote sensory subtype identity. RUNX1 is expressed in TrkA^+^ nociceptive neurons, supporting their differentiation. RUNX3 promotes the segregation of a transient population co-expressing TrkB and TrkC into the TrkC^+^ proprioceptive and TrkB^+^ mechanoreceptive populations by repressing TrkB, both directly and indirectly, while maintaining TrkC expression. Cells that maintain high levels of expression of RUNX3 keep TrkC and become proprioceptors, whereas cells that reduce or lose RUNX3 expression become TrkB^+^/TrkC^+^ or TrkB^+^ mechanoreceptors. Interestingly, RUNX3^−/−^ mice have loss of proprioceptive neurons and severe limb ataxia, features also present in FRDA.

Here we report the use of a combination of small molecules and trophic factors to modulate the pathways involved in the differentiation of DRGs to obtain proprioceptor-enriched primary sensory neuronal cultures from human iPSCs. Proprioceptors can be purified from these cultures by fluorescence-activated cell sorting (FACS). We show that it is possible to reproduce with fidelity the timing of expression of different factors involved in sensory neurogenesis, allowing the study of different stages of the differentiation process and to localize specific differences between healthy controls and different models of diseases affecting the proprioceptive lineage.

## Materials and Methods

### Induced pluripotent stem cell culture

iPSCs were obtained by reprogramming from human fibroblasts from two healthy controls (HEL46.11, HEL24.3) and three FRDA patients (HEL135.2, ULBi004FA4, ULBi005FA1). Participating individuals provided written informed consent and underwent a skin biopsy according to a protocol approved by the institutional Ethics Committee of Erasme Hospital (P2008.313) entitled “Generation of Cellular Models of Neurological Diseases”.

iPSCs were cultured under feeder-free conditions, in Essential 8 Medium (E8, Thermo Fisher Scientific, Cat. No.: A1517001) on Matrigel-coated tissue culture plates (Corning, Cat. No. 356231; 0.05 mg/ml Matrigel solution in DMEM/F12 medium). Cells were fed daily and passaged every 3 days using 0.5 mM EDTA.

### Differentiation of iPSCs into proprioceptor-enriched primary sensory neuron cultures

We modified the dual-SMAD inhibition/WNT activation protocol ^23^ to differentiate primary sensory neurons from human iPSCs to favor differentiation into TrkC^+^ proprioceptors rather than TrkA^+^ nociceptors as in the original report.

iPSCs were plated as single cells on Matrigel treated dishes (0.5 mg/ml Matrigel solution in DMEM/F12 medium) at a density of around 20.000 cells/cm^2^ in E8 medium supplemented with 10 µM ROCK Inhibitor (Y-27632 dihydrochloride, Sigma, Cat. Y0503). The day after, the spent medium was replaced with fresh E8 without Y-27632, and cells were allowed to proliferate until 60-80% confluency (usually reached in 48 hours after seeding). To initiate sensory neuron differentiation, medium was replaced with Essential 6 Medium (E6 Medium, Thermo, Cat. A1516401) supplemented with 100 nM LDN193189 and 10 μM SB431542 (Day 1). E6 medium is equal to E8 minus bFGF and TGFβ proteins, which inhibit differentiation. LDN193189 and SB431542 were added until Day 5 of differentiation. Starting on Day 2, other three small molecules were added: 3 μM CHIR99021 was added from Day 2 to Day 7, while 10 μM DAPT and 9 μM SU5402 where added until Day 8. All inhibitor factors were purchased from STEMCELL Technologies. Cells were fed daily and medium was gradually switched from E6 medium to N2-A medium, starting on Day 4, according to the following schedule: Day 4-5 (75% E6, 25% N2-A), Day 6 (50% E6, 50% N2-A), Day 7-8 (25% E6, 75% N2-A). N2-A medium consists of Neurobasal-A medium (Thermo, Cat. 10888022), supplemented with 1% N2 (100x, Thermo, Cat. 17502001) and 1% GlutaMAX (100x, Thermo, Cat. 35050061). On Day 9, cells were passaged on new Matrigel coated plates to promote maturation and enhance neural survival. At this stage, cells used to be weakly attached to the substrate, so it was possible to gently dissociate them mechanically, without the need of Accutase or other enzymatic solutions, with minimum cellular stress and high survival rate after passaging. The spent medium was removed, cells were rapidly washed once in PBS and fresh medium was added directly to the plate. Cells were mechanically dissociated by gently dispensing the medium a few times against the culture surface and directly transferred into the new plate. It is important for cells to be re-plated at a high confluency to enhance long term survival. The splitting ratio should be adjusted for each line, according to the survival and proliferative rate observed in the first part of the protocol. Starting on Day 9, cells were fed in N2-B medium, consisting of Neurobasal-A medium supplemented with 1% N2, 1% B27 (Thermo, Cat. 17504001), 1% GlutaMAX, 1% MEM Non-Essential Amino Acids (Thermo, Cat. 11140050) and 0.1% β-mercaptoethanol. 40 ng/ml NT3 and 5 ng/ml BDNF were added from this point on, while 5 ng/ml NGF and Glial-Derived Neurotrophic Factor (GDNF) were added until Day 10. Neurotrophic factors were obtained from STEMCELL Technologies. Cells were fed by replacing 75% of medium containing neurotrophins every 2 days. Addition of neurotrophic factors stalls neural proliferation and stimulates a rapid maturation towards a pseudo-unipolar phenotype. The differentiation protocol is summarized in Figure 1.

**Figure 1.**
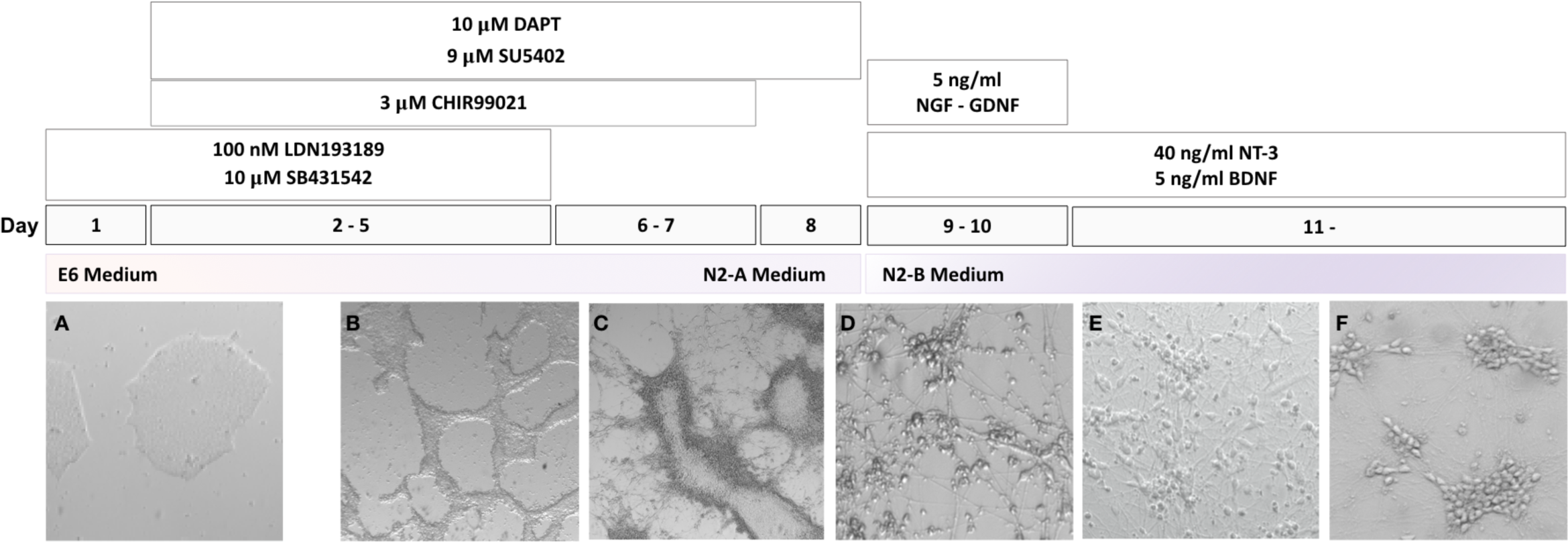
Upper panels: scheme of the differentiation protocol. Lower panels: bright field images of the cultures at each stage of differentiation: **A**. Typical morphology of iPSCs colonies. **B**. Tubular aggregates of neural crest precursors cells. **C**. Initial migration of neural precursors from tubular aggregates. **D**. Neuronal cells migrated out of tubular aggregates, starting to form cellular clusters. **E**. Sensory neuron clusters at low confluency, containing sensory neurons at different stages of differentiation. **F**. Sensory neurons at higher confluency forming large clusters with tight connections.

### Quantitative Real-Time PCR

Cells were harvested at day 2 (first days of differentiation), day 6 (tubular structures), day 8 (migrating cells), day 9 (passage into new plate), and after 3, 5, 7 and 12 days of treatment with neurotrophic factors and processed for total RNA extraction using the RNeasy Mini RNA Kit (Qiagen), following manufacturer’s instructions. Quality of RNA was examined on NanoDrop ND-1000 Spectrophotometer (Isogen) with a A_260/280_ ratio of around 2.0.

Up to 300 ng of total RNA was reverse transcribed using M-MLV Reverse Transcriptase (Thermo, Cat. 28025013) and random primers for cDNA synthesis, according to manufacturer’s protocol. Gene expression was quantified by quantitative reverse transcription PCR (RT-qPCR) on a 7500 Fast RT-PCR System (Applied Biosystem) using Power SYBR Green Master Mix (Thermo, Cat. 4367659). Relative RNA levels were analyzed by the threshold cycle (2^−ΔΔCT^) method normalized to GAPDH expression in iPSCs cells. Expression values are log_2_ of the fold change, with error bars representing standard deviation of mean (SD). Primer sequences are detailed in Supplemental Table 1.

### Immunofluorescence (IF)

Cells grown on Matrigel-coated glass coverslips were fixed in 3.6% paraformaldehyde (PFA, Sigma) for 10 minutes at room temperature and carefully washed twice in PBS after incubation. Membranes were permeabilized with ice cold 0.1% Triton X-100 in PBS for 10 minutes. After washing twice in PBS, non-specific binding sites were blocked with 10% Normal Donkey Serum (NDS, Abcam, Cat. ab7475) for 30 minutes at room temperature. Cells were then incubated with primary antibodies in 5% NDS overnight at 4°C. Following three 5-minute washes in PBS, cells were incubated with secondary antibodies in 5% NDS for 1 hour at room temperature, protected from light. Nuclei were stained with Hoechst 33342 (1 μg/ml, Thermo, Cat. H3750) for 10 minutes at room temperature, protected from light. Samples were then mounted on microscope slides using FluorSave Reagent (Calbiochem, Cat. 345789). Images were acquired using ZEISS Axio Zoom.V16 Microscope or ZEISS AxioImager Z1 (Zeiss, Oberkochen, Germany), and processed using Zeiss ZEN 2.6 Blue Microscopy Software.

Primary antibodies used for immunofluorescence analysis included: SOX10, (1:50, R&D, Cat. MAB2864), Brn3a (1:500, Millipore, Cat. AB5945), Peripherin (1:500, Abcam, Cat. ab99942), Tubulin β-III (1:700, BioLegend, Cat. 801201), TrkA, (1:250, Abcam, Cat. ab76291), TrkB (1:250, Abcam, Cat. ab18987), TrkC (1:250, Abcam, Cat. ab43078 and 1:50, Santa Cruz, Cat. WW6), Pan Trk (1:250, Abcam Cat. ab181560), Parvalbumin (1:250, Abcam Cat. ab11427), Runx3 (1:200, Abcam, Cat. ab135248). Secondary antibodies included: Donkey Anti-Mouse IgG Alexa Fluor 594 (1:1000, Abcam, Cat. ab150118) and Donkey Anti-Rabbit IgG Alexa Fluor 488 (1:1000, Abcam, Cat. ab150073).

During electrophysiological recordings, neurons were filled with biocytin, directly included in the pipette internal solution at a concentration of 0.5% in order to realize a double-staining between biocytin and TrkC to confirm the identity and stage of maturation of patched neurons. Following patch clamp experiments, culture slides were fixed in 3.6% PFA, permeabilized with 0.1% Triton X-100 and blocked with 10% NDS, as previously described. They were incubated overnight at 4°C with anti-TrkC primary antibody in 5% NDS. The day after, following three five-minute washes in PBS, cells were incubated with anti-rabbit secondary antibody (1:1000) and streptavidin-NL557 (1:2000) in 5% NDS for 2 hours at room temperature, protected from light. Culture slides were washed three times in PBS and mounted with FluorSave Reagent for imaging. Z-stack sections were acquired using ZEISS Axio Zoom.V16 Microscope and processed with Zeiss ZEN 2.6 Blue Microscopy Software.

### Electrophysiology

For electrophysiological recordings, individual culture slides from control lines at different stages of maturation (day 7, 10, 12 of treatment with neurotrophic factors) were transferred to a thermoregulated (30-32°C) chamber, maintained immersed and continuously superfused at a rate of 1,5-2 ml/min with oxygenated artificial cerebrospinal fluid (aCSF) containing (in mM): NaCl 127, KCl 2.5, NaH_2_PO_4_ 1.25, MgCl_2_ 1, NaHCO_3_ 26, D-glucose 10, CaCl_2_ 2, bubbled with 95% O_2_ and 5% CO_2_ at a pH of 7.3 (300-316 mOsm). Patch clamp experiments were performed in whole cell configuration on individual neurons, identified with a 63x water immersion objective from Zeiss Axioskop microscope (Axioskop 2FS Plus; 140 Zeiss, Oberkochen, Germany) with an infrared CCD camera (X-ST70CE, Hamamatsu Photonics KK, Hamamatsu, Japan). Cells were selected based on their size and morphology; as proprioceptive neurons are reported to be among the largest sensory neurons in DRGs, only large and pseudo-unipolar neurons were patched.

Borosilicate-glass patch electrodes [4-6MΩ, (Hilgenberg GmbH, Malsfeld, Germany)] were filled with a solution containing biocytin 0.5% (Sigma-Aldrich, Cat. B4261) and (in mM): KMeSO_4_ 125, KCl 12, CaCl_2_ 0.022, MgCl_2_ 4, HEPES 10, EGTA 0.1, Na_2_-phosphocreatine 5, Mg_2_-ATP 4, Na_2_-GTP 0.5 (pH of 7.2, 292 mOsm). Currents were recorded using an EPC-10 patch clamp amplifier (HEKA, Lambrecht, Germany) and PatchMaster acquisition software (HEKA).

Cells were first recorded in voltage-clamp mode at holding potential of −60 mV with a gain of 2 mV/pA and low-pass filtered at 2.9 kHz. Signals were sampled at 20 kHz. Passive membrane properties (Capacitance (Cm, pF), Membrane resistance (Rm, MΩ), Membrane time constant (τ, ms)) and series resistances were extracted from current traces recorded in response to a hyperpolarizing voltage pulse (200 ms) of −10 mV from holding potential. Ten sweeps were averaged to remove noise.

Then, in current-clamp mode, cell excitability was investigated by setting the resting membrane potential at −60 mV and injecting 1 s depolarizing steps (from 0 to 100 pA in 10 pA increments). Signals were sampled at 10 kHz with a gain of 2 mV/pA. To measure the neuron resting membrane potential (RMP), the potential fluctuations over the duration of the step at 0 pA of injected current were averaged off-line. For cells recorded with spontaneous action potentials (APs), the RMP was measured manually by the value of neuron potential at 0.02s before the first action potential of the step at 0 pA. Series resistance was not compensated during the recordings and membrane potential values were corrected off-line with a liquid junction potential of 6.6 mV. If access resistance exceeded 35 MΩ and changed more of 25% between the beginning and the end of the recording, the neuron was discarded. The series resistance averaged for all recordings is of 27,32±1,331 MΩ.

Analysis of passive properties and excitability of patched neurons were performed with IgorPro 6.3 software (WaveMetrics, Portland, USA) using Patcher’sPower Tools, NeuroMatic plugins and Microsoft Excel software. Results are expressed as mean±SEM.

When possible, confirmation of morphology and identity of recorded neurons was assessed with biocytin-TrkC double immunostaining, as described in the previous section (N=7).

### Flow cytometry

For Flow Cytometry, neurons were harvested after one week of treatment with neurotrophins using StemPro Accutase and gently triturated with a 1000 μl pipette to prepare a single cell suspension. Cell clumps were removed by filtration of suspension through a 70 μm strainer. Cells were counted and spun at 200 g for three minutes. The cell pellet was resuspended at a concentration of 10^7^ cells/ml in cold Flow Cytometry (FC) Staining Buffer 1x (R&D, Cat. FC001) and transferred into a protein low-binding tube. 3×10^5^ cells were transferred into another tube and used as negative control for FACS. Samples were incubated with an Anti-TrkC antibody conjugated with Phycoerythrin (10 µl/10^6^ cells, R&D, Cat. FAB373P) for 1 hour on ice, protected from light. Cells were mixed by gently inverting the tube 3-5 times every 5-10 minutes during incubation. They were then spun at 300 g for 3 minutes at 4°C, washed with 500 μl of FC Staining Buffer 1x and resuspended in 200 μl of the same buffer. TrkC^+^ cells were sorted in a BD FACS Aria II System (BD Bioscience) and directly dissolved in RTL buffer for subsequent RNA extraction and RT-qPCR analysis. Efficiency of sorting was assessed by comparison of Trk receptors expression between sorted cells and original cultures.

## Results

### iPSC differentiation into proprioceptor-enriched primary sensory neuron cultures

Figure 1 shows a summary of the steps of the differentiation protocol and the corresponding morphology of cell cultures by bright field microscopy. The process of iPSC differentiation in culture recapitulated the *in vivo* DRG development. Initially, iPSCs proliferated in a uniform confluent layer, without any significant morphological changes. Between day 5 and 6, cells rapidly formed regular tubular aggregates where they were tightly connected. On day 7, cells with the morphology of neural precursors started migrating out of the tubular aggregates, which rapidly disappeared in the following two days, forming a uniform layer. At this stage (day 9), cells were passed into a new plate and allowed to mature in the presence of neurotrophic factors. In this second phase, cells showed rapid morphological changes, from bipolar, to bell-shaped to pseudo-unipolar neurons. Such phenotypical maturation was coupled with the organization of neurons in clusters and with the generation of a tight network of neurites.

The timing and efficiency of the process were investigated by qRT-PCR and by IF analysis of the expression of transcription factors involved at different stages of sensory neurogenesis and of other markers of pluripotent cells, neural crest cells, developing and mature sensory neurons. At the beginning of the differentiation process cells rapidly lost the expression of the pluripotent marker OCT4, retaining the marker for multipotent neural stem cells SOX2 until day 9 (Figure 2A-B). At the same time the human neuroectoderm marker PAX6 was transiently induced up to 4-fold iPSC level, indicating that the treatment mostly generated cells of the sensory lineage (Figure 2C). Expression of the neural crest marker SOX10 peaked at day 6 (4-fold compared to iPSC level) then rapidly declined, returning to iPSC level or lower at day 9, indicating a transient transition through a neural crest phenotype (Figure 2D). BRN3A, marker of developing and mature sensory neurons, showed a strong induction (3-5×10^3^-fold), plateauing around day 8-9 when SOX10 was declining (Figure 2E). Figure 3A-C shows co-expression of BRN3A and SOX10 in tubular structures at day 6, Figure 3D-F shows persisting BRN3A expression in differentiated sensory neurons labeled with the specific neuronal marker TUBB3. Neurogenin 1 and Neurogenin 2 expression peaked between day 6 and 9, with a robust 3-5×10^3^-fold increase compared to iPSC levels, without clearly showing the two distinct waves occurring *in vivo* (Figure 2F-G). Their expression declined when neurotrophic factors were added, leading to the final sensory neuron differentiation. Addition of neurotrophins and expression of Trk receptors (Figure 4 and 5) marked the final stage of sensory neurogenesis. The brief (day 9-10) exposure to low levels (5 ng/ml) of NGF in the second part of the protocol transiently sustained the differentiation and survival of nociceptors, as also indicated by moderate induction of RUNX1 (Figure 2H). However, nociceptors constituted a minority of the differentiated sensory neurons and they were progressively lost after the removal of NGF from the culture medium, as indicated by a progressive reduction in the expression of TrkA (Figure 4A and 5A-B), which was eventually 10^3^-fold lower in differentiated cultures than at the beginning of final sensory neuron differentiation (day 9). The expression of TrkB continued instead to increase until day 15, then remained stable despite the low concentration of BDNF in the culture (5 ng/ml) (Figure 4B and 5C-D), probably due to its tight association with TrkC. Contrary to the other Trk receptors, TrkC was already robustly expressed in iPSCs, then showed a slight (2-8-fold) decrease in the first part of differentiation. This is consistent with the observation that TrkC is among the earliest markers of sensory neurogenesis *in vivo* and is already expressed by migrating neural crest and neural progenitor cells. TrkC expression increased again in the final differentiation stage, when a high concentration of NT-3 (40 ng/ml) was used, to up to 100-fold its iPSC level (Figure 4C). At this stage about half of cells appeared to express TrkC by IF (Figure 5E-G). P75^NTR^, member of the tumor necrosis factor receptor family, is a low affinity receptor for all neurotrophins expressed on neural crest cells and co-expressed with Trk receptors in sensory neurons. It functionally collaborates with Trk receptors to enhance responses to preferred Trk ligands, to reduce responses to non-preferred ligands, and to facilitate apoptosis resulting from neurotrophin withdrawal. The expression of p75^NTR^ was progressively induced throughout all phases of the treatment and persisted in mature neurons (Figure 4D). The presence of primary sensory neurons was confirmed by strong expression of the general neural marker Tubulin-β-III and the primary sensory neuronal marker Peripherin (Figure 6). Specific markers of proprioceptive neurons that were induced in the final stage of differentiation included the transcription factor RUNX3 (Figure 2I), vGLUT1 and PV (Figure 4 E-F). IF confirmed PV and RUNX3 co-expression (Figure 7). RUNX3 and PV expression was relatively high in iPSCs, then it declined during neural crest specification and increased again in the second part of the protocol during the final specification and maturation of proprioceptive neurons, returning to or exceeding iPSC levels in control cells but not in FRDA cells (Figure 2G and 4F).

**Figure 2.**
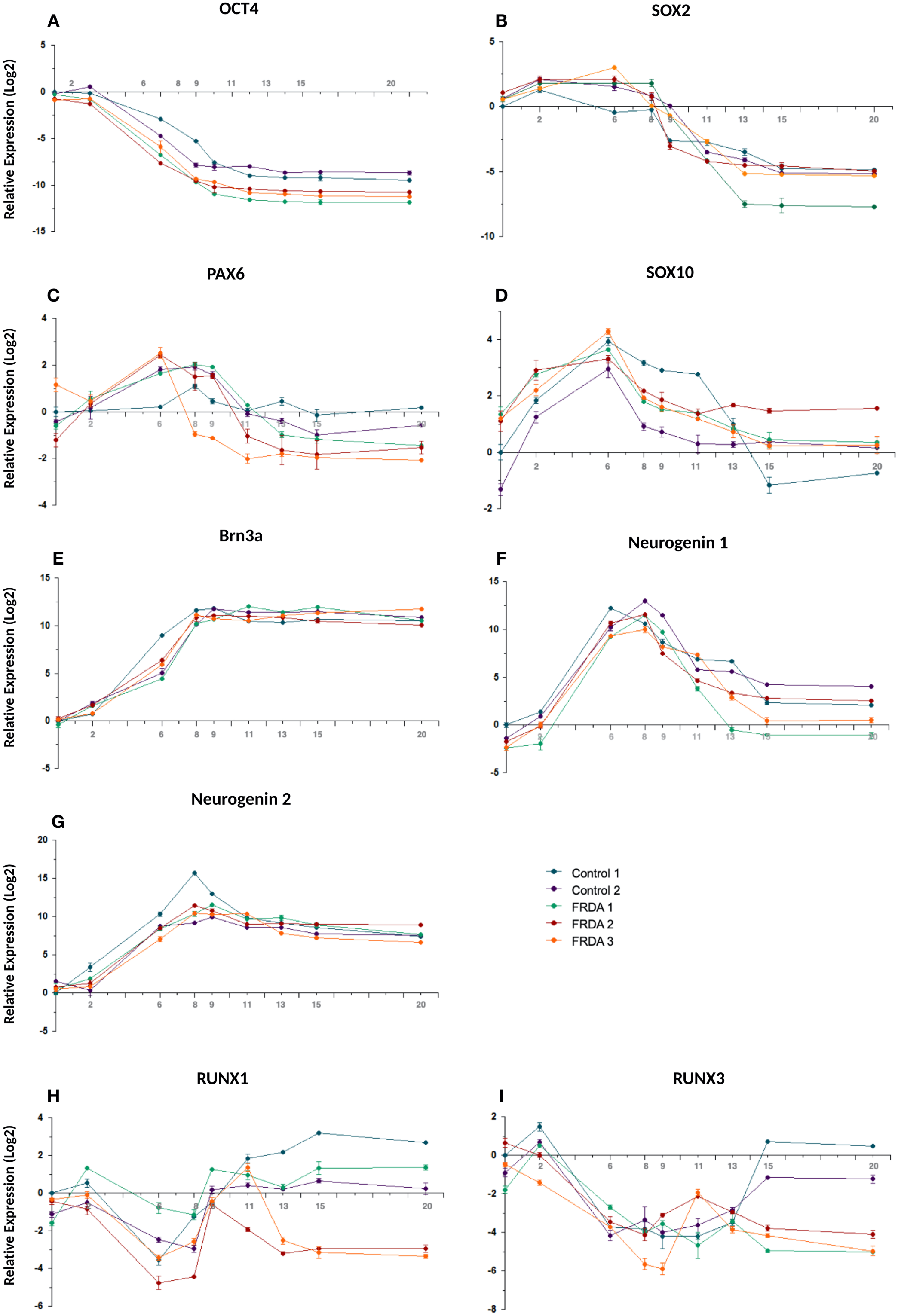
Time course of the expression of nine transcription factors during iPSC differentiation to sensory neurons in two control and three FRDA lines. mRNA levels were determined by quantitative RT-PCR in duplicate experiments.

**Figure 3.**
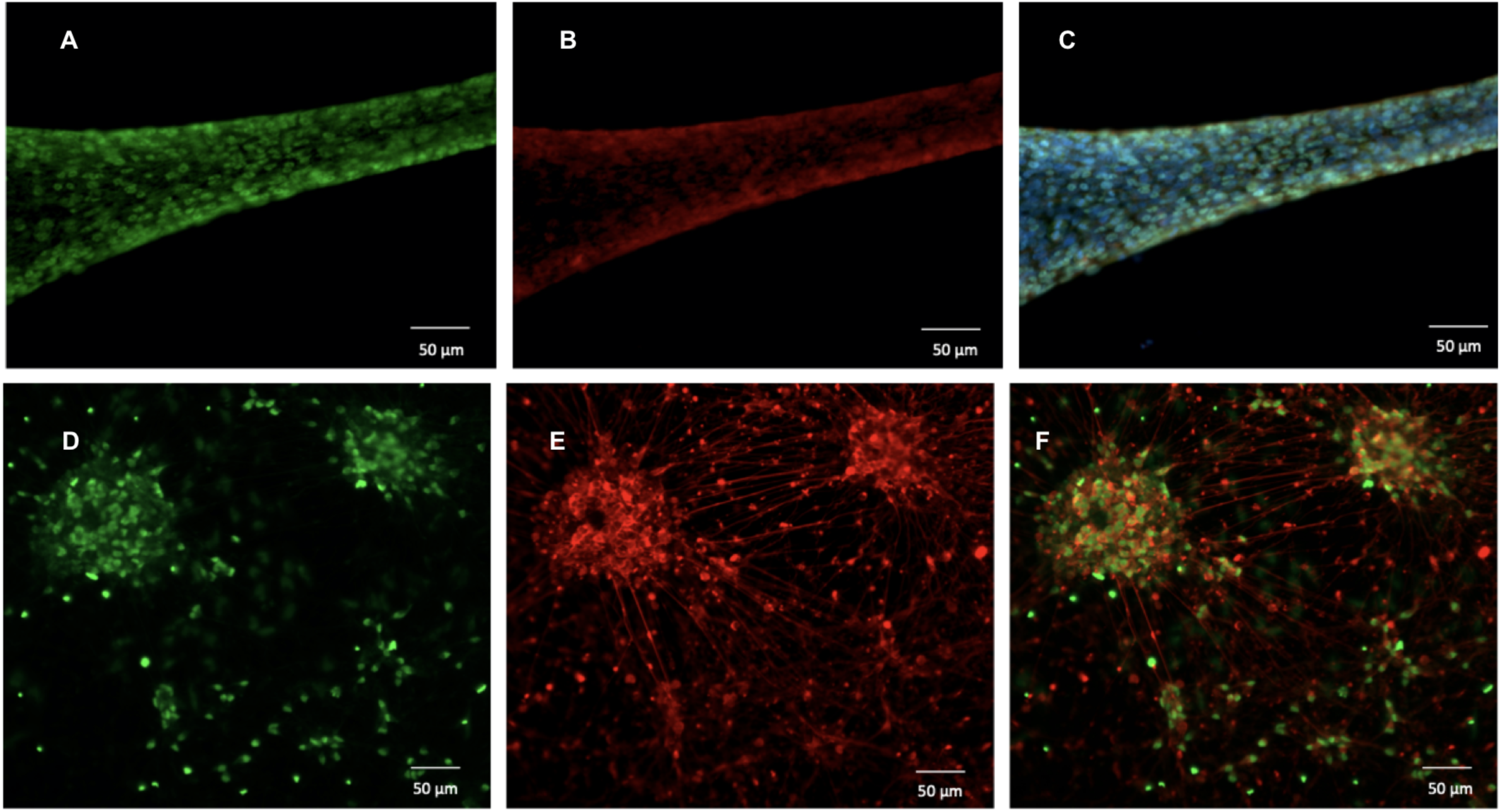
**(A-C)** SOX10 (green, panels A and C) and BRN3A (red, panels B and C) expression by IF in tubular aggregates of neural precursors at day 5. Nuclei are shown in blue (DAPI) in panel C. **(D-F)** BRN3A (green, panels D and F) TUBB3 (red, panels E and F) expression by IF in differentiated sensory neurons at day 15. Nuclei are shown in blue (DAPI) in panel F. Images were acquired with ZEISS Axio Zoom.V16 microscope, magnification: 179x.

**Figure 4.**
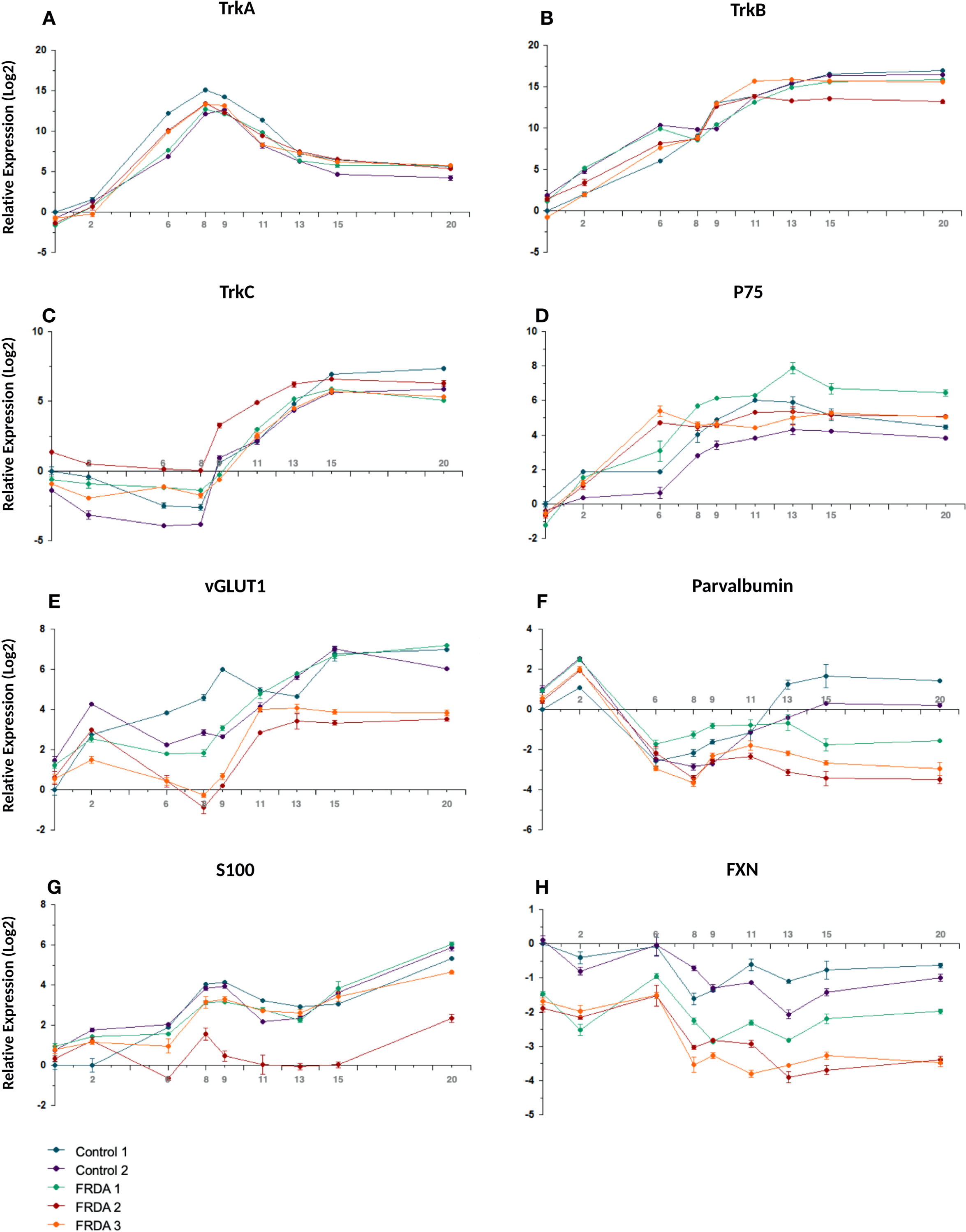
Time course of log2 expression of Trk receptors, p75, vGLUT1, PV, S100 and FXN during iPSC differentiation to sensory neurons in two control and three FRDA lines. mRNA levels were determined by quantitative RT-PCR in duplicate experiments and normalized to levels in control iPSCs.

**Figure 5.**
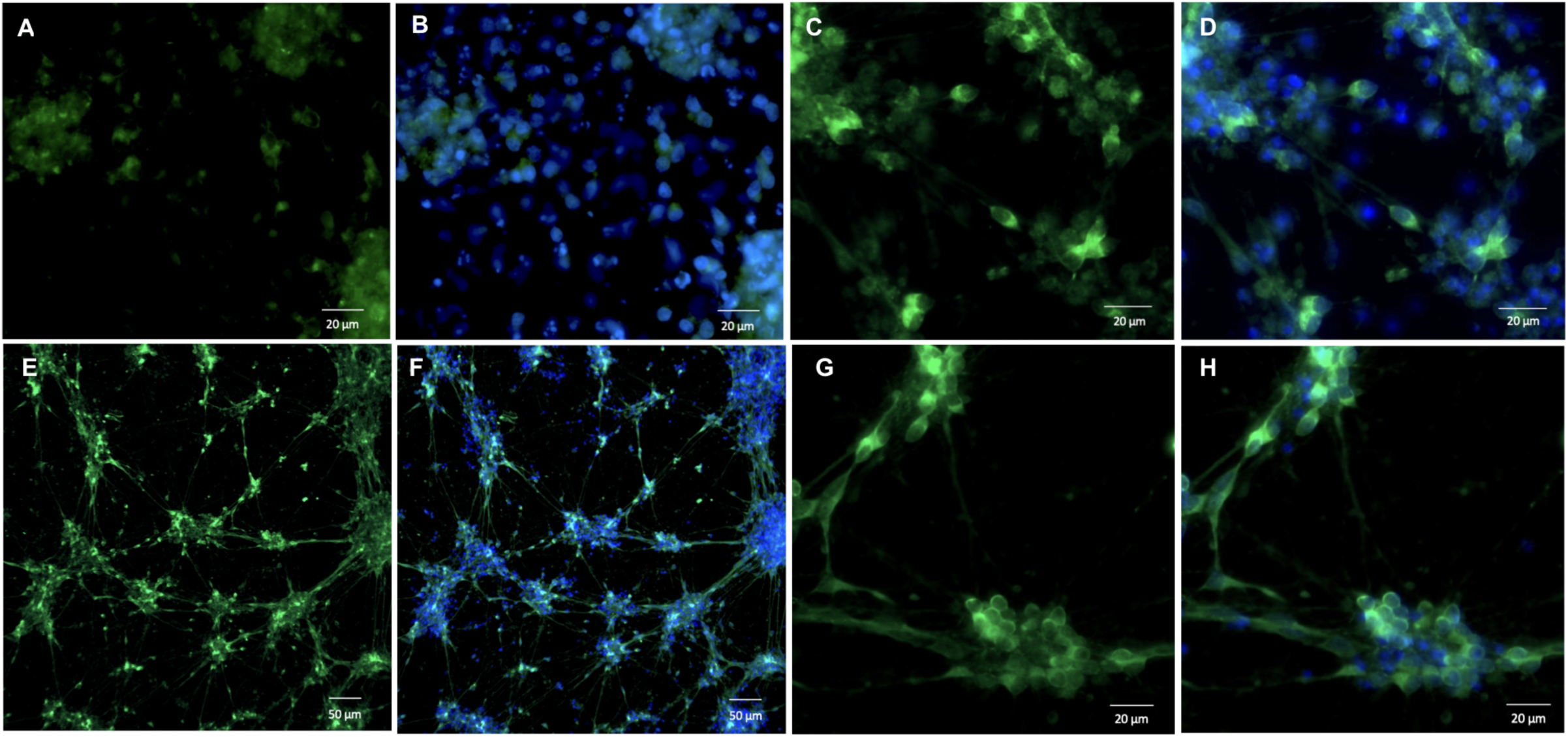
**(A, B)** TrkA; **(C, D)** TrkB; **(E-G)** TrkC expression by IF in differentiated sensory neurons at day 15. Staining for all antibodies is in green. Nuclei are shown in blue (DAPI) in panels B, D, F, H. Images were acquired with ZEISS Axio Zoom.V16 microscope. Panels A, B, C, D, G, H magnification: 412x, panels E, F magnification: 179x.

**Figure 6.**
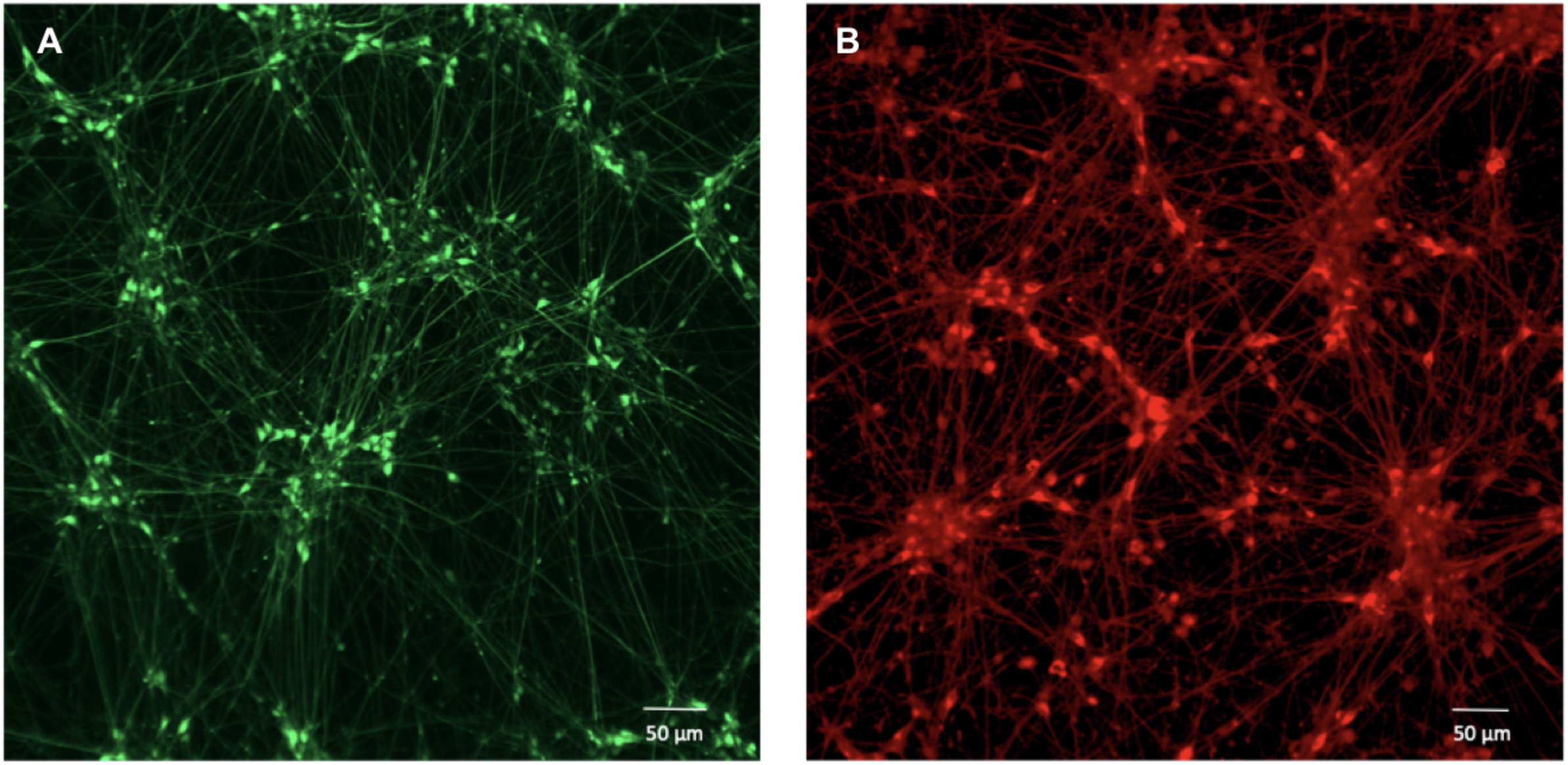
**(A)** IF image showing peripherin (green), expression in differentiated sensory neurons at day 15. **(B)** IF image showing TUBB3 (red) expression in differentiated sensory neurons at day 15. Images were acquired with ZEISS Axio Zoom.V16 microscope, magnification: 179x.

**Figure 7.**
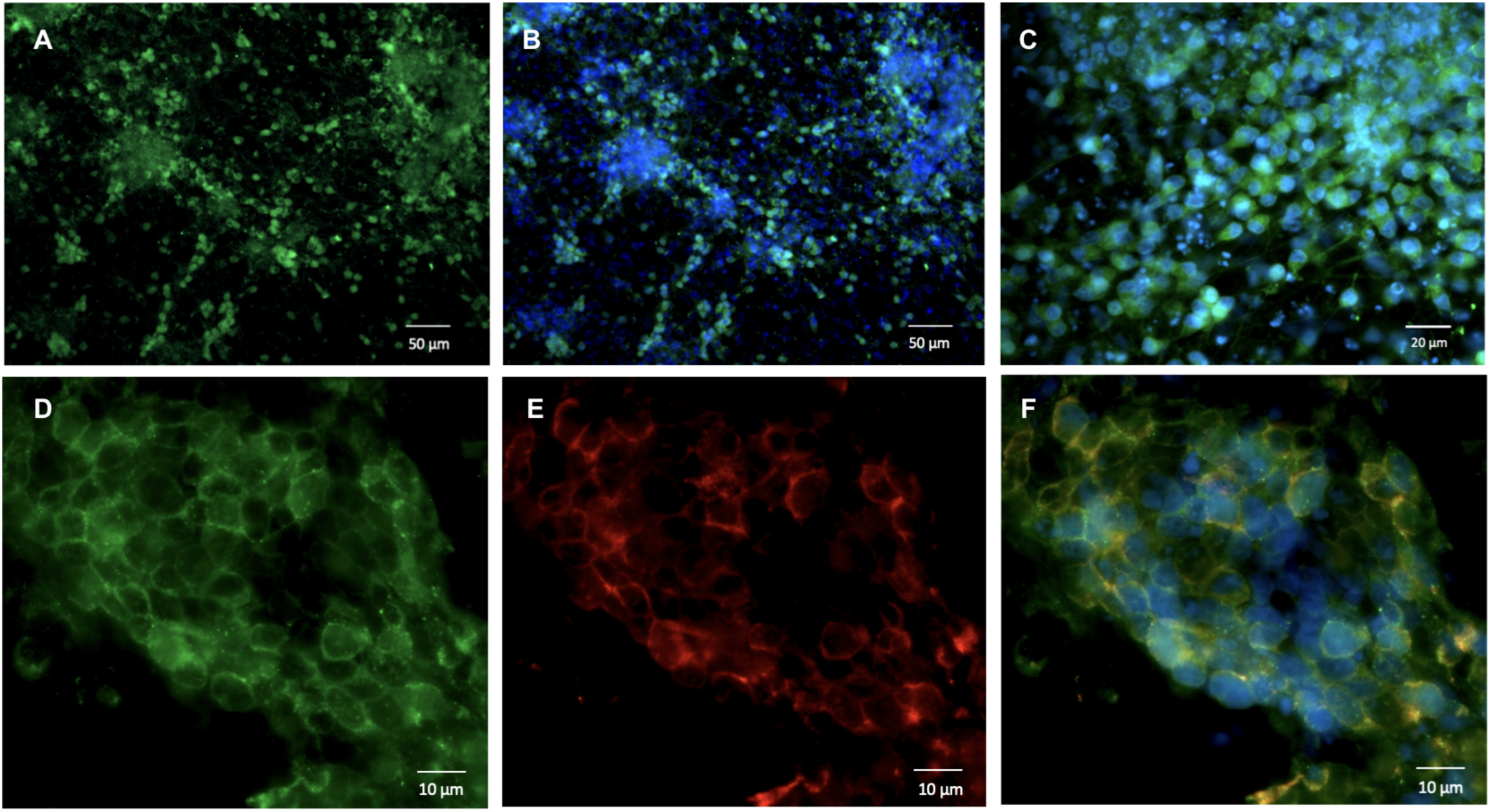
**(A-C)** IF images showing PV (green) expression in differentiated sensory neurons at day 15. Nuclei are shown in blue (DAPI) in panels B and C. **(D-F)** IF images showing PV (green, panels D, F) and RUNX3 (red, panels E, F) expression in differentiated sensory neurons at day 15. Nuclei are shown in blue (DAPI) in panel F. Images were acquired with ZEISS Axio Zoom.V16 microscope Panels A, B magnification: 179x, panel C magnification: 412x, panels D-F magnification: 895x.

S100, a marker for glia and satellite cells in DRGs, showed progressive induction with a transient small dip at the beginning of the final sensory neuronal differentiation (Figure 4G). FXN expression was 3-4-fold lower in FRDA iPSCs and throughout the differentiation process. It showed a mild decline in all lines after neural crest induction around day 6 (Figure 4H).

Overall, we could mimic the initial *in vivo* sensory neuron differentiation and DRG development, then strongly enriched for cells expressing proprioceptive neurons markers in the final phase of sensory neuronal type determination. By IF analysis, TrkB^+^ and TrkC^+^ expressing cells mostly formed separate populations with few cells still expressing both receptors. Semi-quantitative assessment of PV^+^, and RUNX3^+^ cells in IF images (Figure 7), in addition to TrkC (Figure 5E-G), indicated that they represented about half of all cells, a 4-5-fold enrichment compared to the physiological composition of DRG neurons.

### FACS purification of proprioceptors

We FACS-purified TrkC^+^ neurons from differentiated iPSC-derived sensory neuron cultures. Though we labeled living cells with an antibody against a surface antigen, we did not put them back in culture because of the damage caused by the procedure that necessarily severs neurites. Differentiated primary sensory neurons were collected at day 15 and stained with an anti-TrkC primary antibody conjugated with Phycoerythrin (PE) for FACS purification. FACS data showed that the percentage of sorted TrkC^+^ cells was around 34% (Figure 8A-B). This is slightly less than estimated by IF, probably because only cells with the highest expression of TrkC were sorted, most likely to be proprioceptors. Furthermore, Trk receptors are rapidly internalized in stress conditions. The identity of sorted cells was confirmed by quantitative RT-PCR, showing a 6-fold enrichment for TrkC mRNA in sorted cells compared to the original cultures (Figure 8C). Of notice, TrkB expression was only marginally increased in purified cells, indicating that only a limited number were TrkB^+^/TrkC^+^ double positives. Interestingly, FXN expression was 50% higher in sorted cells, in agreement with higher expression in proprioceptors (Figure 8C).

**Figure 8.**
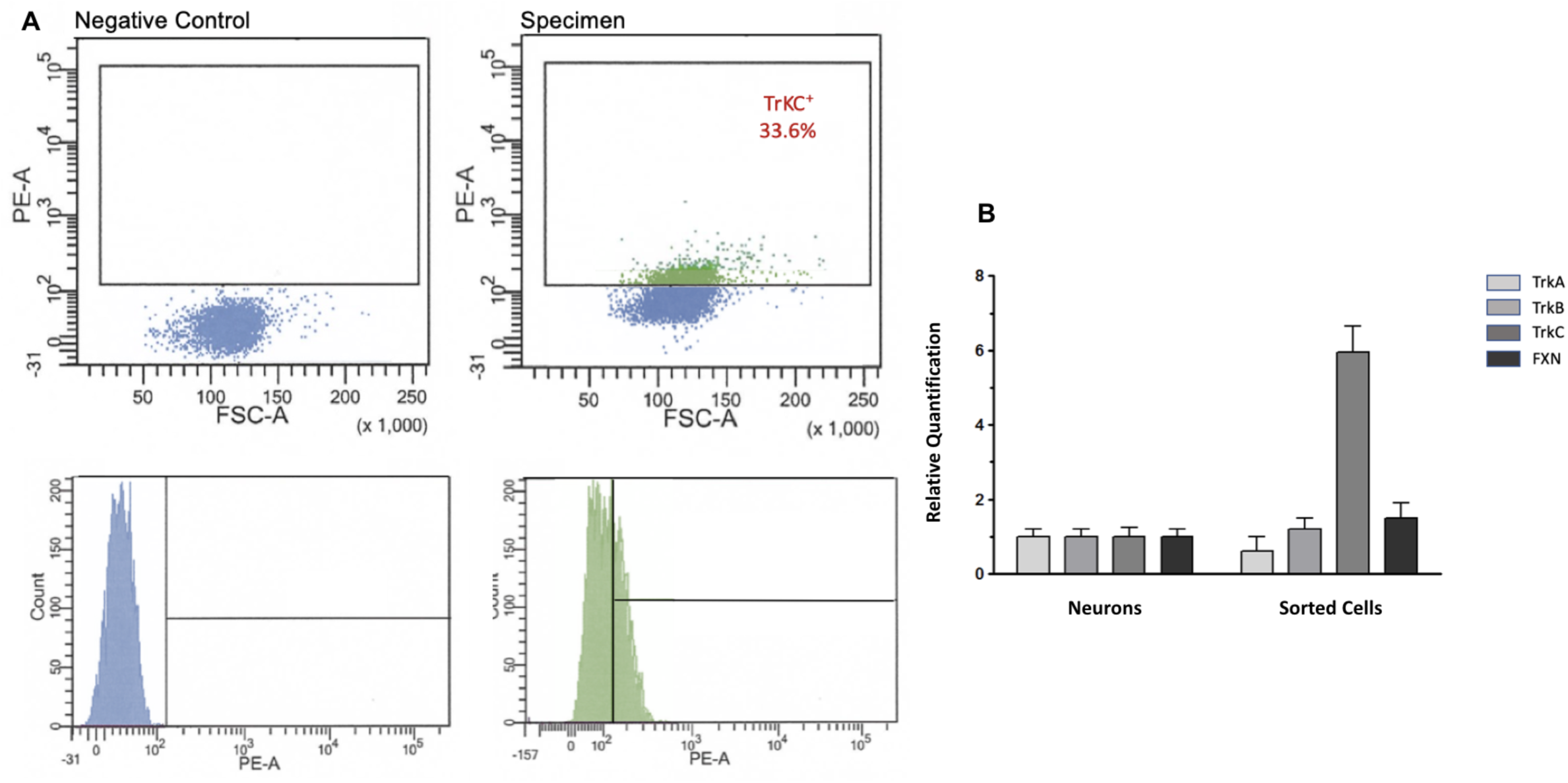
**(A)** FACS purification of TrkC+ iPSC-derived primary sensory neurons (negative control on the left) from iPSC-derived peripheral sensory neurons. **(B)** Fold-difference of Trk receptor expression by quantitative RT-PCR analysis in sorted cells vs. starting sensory neurons cultures.

### Functional characterization of differentiated proprioceptive neurons

Whole-cell patch-clamp recordings were performed to characterize the functional maturity of iPSC-derived neurons. A time course analysis of the electrophysiological excitability of these neurons in current clamp configuration (depolarizing current injection steps from 0 pA to 100 pA) was first performed in order to assess the timing of functional maturation. This analysis started at day 15, when cells are already organized in clusters and show a mature morphological phenotype (Figure 9A), and was pursued at day 18 and 20. The majority of neurons patched at the first stage showed only one AP at the beginning of the depolarizing current step, with just a few of them being able to generate a short train of low amplitude APs with rapid accommodation for increasing currents. Three days after, neurons exhibited an improved spiking profile, but were still characterized by a rapid accommodation in the majority of cases (Figure 9B).

**Figure 9.**
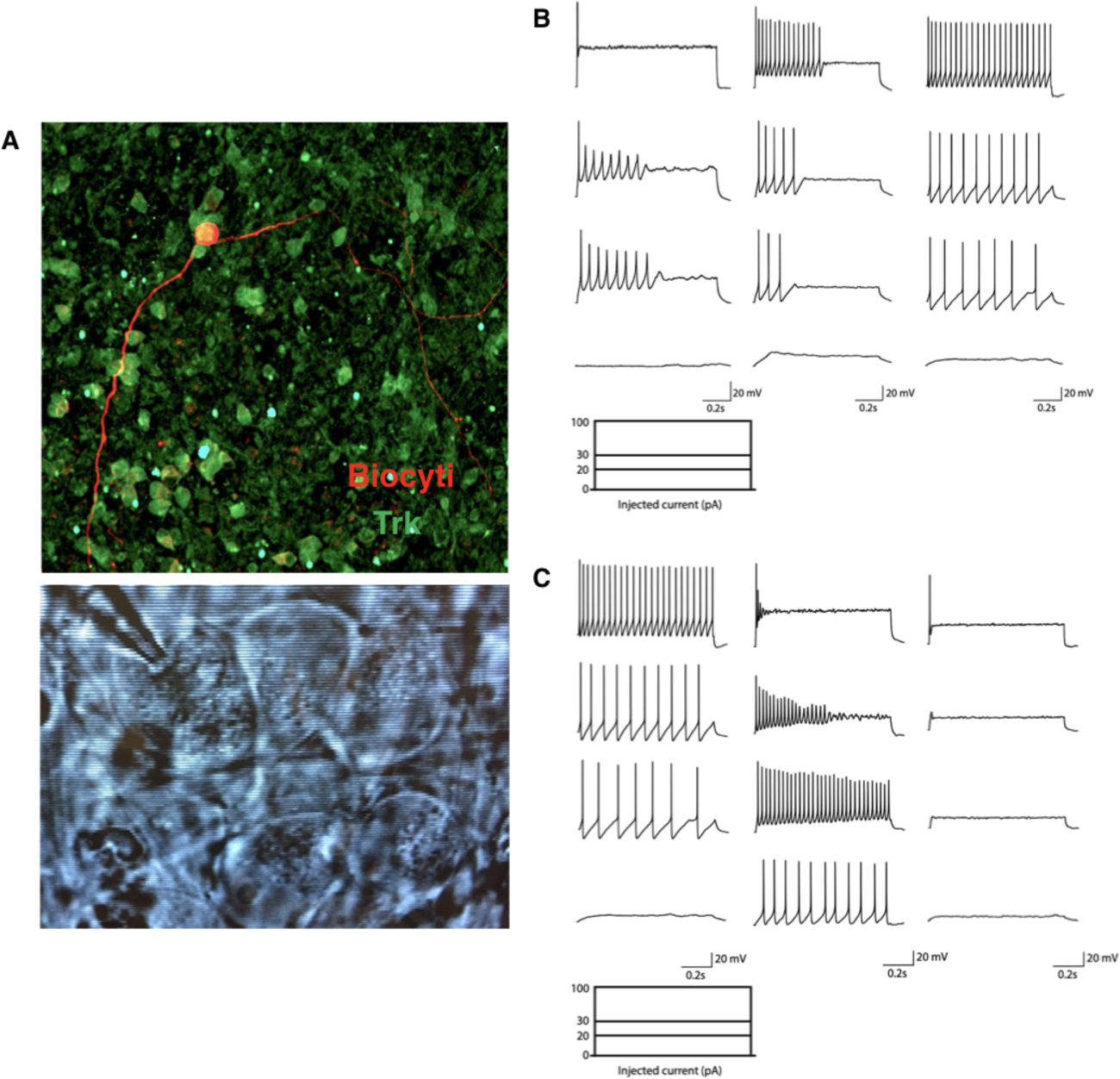
**(A)** Time course analysis of electrophysiological excitability of differentiated neurons. Representative traces of current-clamp recordings in cells at day 7 (left) 10 (middle) or 12 (right) of treatment with neurotrophic factors, in response to depolarizing current injection steps at 0 pA (lower trace) 20, 30 and 100 pA (higher trace). **(B)** (Up) Representative image of biocytin-filled TrkC^+^ neuron. (Down) Representative image of neurons identified with a 63x water immersion objective and an infrared CCD camera during electrophysiological recording. They showed the typical morphology of large sensory pseudo-unipolar neurons. **(C)** Representative traces of current-clamp recordings in cells at day 12 of treatment with neurotrophic factors, in response to 0 (lower traces) 20, 30 and 100 pA (higher traces) of depolarizing current injection steps. Three different responses were observed: regular APs firing with firing frequency increasing for higher current injections (left); spontaneous activity with rapid accommodation following increasing step current injections (middle); single AP in response to higher injecting currents (right).

Finally, at day 20 (N=15), it was possible to observe three different firing behaviors in the cultures (Figure 9C): some neurons were capable of firing and sustaining regular APs in response to depolarizing current injections, with firing frequency increasing in response to higher currents; some neurons exhibited spontaneous AP firing but rapid accommodation following increasing step current injection; finally, some neurons were able to generate a single AP at the beginning of the depolarizing step in response to higher injecting currents. On average, recorded neurons exhibited a resting membrane potential of −62,75± 0,7493 mV.

Intrinsic passive properties of differentiated neurons at day 20 were obtained in voltage clamp configuration recording. Membrane resistance was 1963 ± 385.0 MΩ. The average capacitance was 29,65 ±1,764 pF and the membrane time constant τ was 51.29 ± 6.35 ms.

### Cellular phenotype of FRDA primary sensory neurons

All three FRDA iPSC lines and differentiated sensory neurons maintained GAA repeat expansions in the *FXN* gene (not shown) and had repressed FXN expression (Figure 4H). They could differentiate similarly to control cells and give rise to very similar mixed cultures of sensory neurons. However, some differences emerged in the final differentiation stage. First, FRDA differentiated sensory neurons had lower viability and shorter survival, which rarely went beyond day 15. The few FRDA neurons that survived beyond day 15, however, had similar neurophysiological properties as control neurons. In all three FRDA lines loss of viability was accompanied by a decrease in the expression of PV and RUNX3, both proprioceptor-specific markers, starting on day 13 (Figure 2G and 4F), when the final segregation of proprioceptive neurons from TrkB^+^/TrkC^+^ lineage was expected to be completed. Differences for other markers were less consistent. In particular, Trk receptors were similarly expressed by control and FRDA lines and only two FRDA lines showed lower levels of the proprioceptor marker vGLUT1.

## Discussion

Numerous protocols have been developed for the generation of primary sensory neurons from pluripotent stem cells. However, they either do not enrich for a specific sensory neuron subtype or they focus on the generation of nociceptive neurons for the study of pain-related disorders. We developed a method to differentiate iPSCs into a primary sensory neuron population showing a substantial enrichment for proprioceptors, which can be FACS-purified for experiments as transcriptome analysis, including plate-based deep single-cell profiling^24^.

The cellular behavior and the pathways activated in the *in vitro* differentiation process resemble those observed during DRG development *in vivo*. We could robustly induce the transcription factors involved in sensory neurogenesis, driving the differentiation of iPSCs into cells expressing specific markers of DRG neurons. Furthermore, differentiated neurons could establish mature networks in just three weeks after plating, which is much faster than what is usually observed with iPSC-derived neurons. The specific timing of treatment of iPSCs with inhibitory factors was able to induce the rapid acquisition of a sensory precursor phenotype, while the combination of neurotrophic factors used in the stage of final differentiation and maturation allowed the generation of a high proportion of sensory neurons of the TrkB^+^/TrkC^+^ lineage. Nociceptive neurons depending on NGF for their survival were largely removed from the final culture, as this neurotrophin was only briefly added at low concentration. Some residual expression of TrkA in mature cultures was probably due to the activity of NT-3, which is able to bind, even if with low affinity, also to TrkA and TrkB. We used a high concentration of NT-3 and a low concentration of BDNF to favor final differentiation into TrkC^+^ proprioceptive rather than TrkB^+^ mechanoreceptive neurons. We obtained this way a substantial enrichment for TrkC^+^ cells, which represented at least a third of the final cultures. However, we still obtained a substantial proportion of TrkB^+^ cells. These two classes of sensory neurons derive from a common precursor are share many morphological and functional features. Their segregation depends on the combination of intrinsic and extrinsic pathways, which are very difficult to modulate *in vitro*. Furthermore, some mature neurons continue to co-express TrkB and TrkC. For this reason, we set up a FACS purification procedure to specifically collect cells with high TrkC expression, almost all of which are expected to be proprioceptors, for specific assays like RNAseq. Not aiming to put cell back in culture, FACS can also be performed in fixed cells using specific intracellular markers for proprioceptive neurons, such as PV and RUNX3.

We confirmed that FXN expression is repressed in FRDA cell lines, consistent with levels found in FRDA patients. We also started to address the issue of a possible developmental defect of proprioceptors in FRDA. FRDA sensory neurons showed declining levels of the proprioceptive neuron-specific marker PV and lower levels of the transcription factors RUNX3, essential for proprioceptive neuron differentiation and survival, particularly in later stage cultures. Correspondingly, differentiated FRDA cultures were less viable and often did not survive after day 15.

Though we developed this model because of our interest in translational research in FRDA, it is of obvious interest to investigate many other conditions affecting proprioceptors, both genetic and acquired. The availability of human proprioceptors carrying the causative genetic mutations, or exposed to pathogenic factors as toxins or autoantibodies, allows to directly explore the relevant pathogenic mechanisms better than primary DRG cultures from animal models that often are only approximations for the human condition.

## Acknowledgments

We thank the donors of the cells used for these experiments. This study was partially supported by a grant to MP from the Friedreich Ataxia Research Alliance. CD is supported by a doctoral fellowship from the Belgian national Scientific Research Funds (FNRS).

## Author Information

### Contributions

C.D., M.R., M.C, S.N.S. and M.P. designed the experiments, C.D. and M.C. conducted experiments and analyzed data, C.D., M.R., M.C, S.N.S. and M.P. interpreted the results. C.D. and M.P. drafted the manuscript and prepared the figures. All authors reviewed the manuscript for intellectual content.

