## Supplementary Table 1 for "Induced pluripotent stem cell-derived primary proprioceptive neurons as Friedreich ataxia cell model"

Supplementary materials

**SUPPLEMENTARY TABLE 1**. List of primers used for RT-qPCR.

| **Gene** | **Sequence (5’-3’)** |
| --- | --- |
| **GAPDH** | **F** GGAGCGGAGTCCCTCCAAAAT |
|  | **R** GGCTGTTGTCATACTTCTCATGG |
| **SOX2** | **F** GCCGAGTGGAAACTTTTGTCG |
|  | **R** GGCAGCGTGTACTTATCCTTCT |
| **OCT4** | **F** GTGTTCAGCCAAAAGACCATCT |
|  | **R** GGCCTGCATGAGGGTTTCT |
| **SOX10** | **F** CCTCACAGATCGCCTACACC |
|  | **R** CATATAGGAGAAGGCCGAGTAGA |
| **BRN3A** | **F** GGGCAAGAGCCATCCTTTCAA |
|  | **R** CTGTTCATCGTGTGGTACGTG |
| **NEUROGENIN 1** | **F** GCTCTCTGACCCCAGTAGC |
|  | **R** GCGTTGTGTGGAGCAAGTC |
| **NEUROGENIN 2** | **F** AGGAAGAGGACGTGTTAGTGC |
|  | **R** GCAATCGTGTACCAGACCCAG |
| **PAX6** | **F** TGGGCAGGTATTACGAGACTG |
|  | **R** ACTCCCGCTTATACTGGGCTA |
| **TRKA** | **F** GCTGGCTCTTCAATGGCTC |
|  | **R** GTGTAGTTGCCGTTGTTGACG |
| **TRKB** | **F** TCGTGGCATTTCCGAGATTGG |
|  | **R** TCGTCAGTTTGTTTCGGGTAAA |
| **TRKC** | **F** CTTTGCCCAGCCAAGTGTAGT |
|  | **R** CGTGATGTTGATACTGGCGTT |
| **P75NTR** | **F** CCTACGGCTACTACCAGGATG |
|  | **R** CACACGGTGTTCTGCTTGT |
| **vGLUT1** | **F** CAGAGTTTTCGGCTTTGCTATTG |
|  | **R** GCGACTCCGTTCTAAGGGTG |
| **PARVALBUMIN** | **F** GCTGAACGCTGAGGACATCAA |
|  | **R** ACATCATCCGCACTCTTTTTCTT |
| **RUNX1** | **F** TGAGCTGAGAAATGCTACCGC |
|  | **R** ACTTCGACCGACAAACCTGAG |
| **RUNX3** | **F** AGGCAATGACGAGAACTACTCC |
|  | **R** CGAAGGTCGTTGAACCTGG |
| **S100** | **F** TGGCCCTCATCGACGTTTTC |
|  | **R** ATGTTCAAAGAACTCGTGGCA |
| **FXN** | **F** CAGAGGAAACGCTGGACTCT |
|  | **R** AGCCAGATTTGCTTTGG |
